# Quantitative proteomics identifies tumour matrisome signatures in patients with non-small cell lung cancer

**DOI:** 10.1101/2022.09.29.510064

**Authors:** Helen F. Titmarsh, Alex von Kriegsheim, Jimi C. Wills, Richard A. O’Connor, Kevin Dhaliwal, Margaret C. Frame, Samuel B. Pattle, David A. Dorward, Adam Byron, Ahsan R. Akram

## Abstract

The composition and remodelling of the extracellular matrix (ECM) are important factors in the development and progression of cancers, and the ECM is implicated in promoting tumour growth and restricting anti-tumour therapies through multiple mechanisms. The characterisation of differences in ECM composition between normal and diseased tissues may aid in identifying novel diagnostic markers, prognostic indicators and therapeutic targets for drug development. Using tissue from non-small cell lung cancer (NSCLC) patients undergoing curative intent surgery, we characterised quantitative tumour-specific ECM proteome signatures by mass spectrometry, identifying 161 matrisome proteins differentially regulated between tumour tissue and nearby non-malignant lung tissue. We defined a collagen hydroxylation functional protein network that is enriched in the lung tumour microenvironment. We validated two novel putative extracellular markers of NSCLC, the collagen cross-linking enzyme peroxidasin and a disintegrin and metalloproteinase with thrombospondin motifs 16 (ADAMTS16), for discrimination of malignant and non-malignant lung tissue. These proteins were up-regulated in lung tumour samples, and high *PXDN* and *ADAMTS16* gene expression was associated with shorter survival of lung adenocarcinoma and squamous cell carcinoma patients, respectively. These data reveal tumour matrisome signatures in human NSCLC.

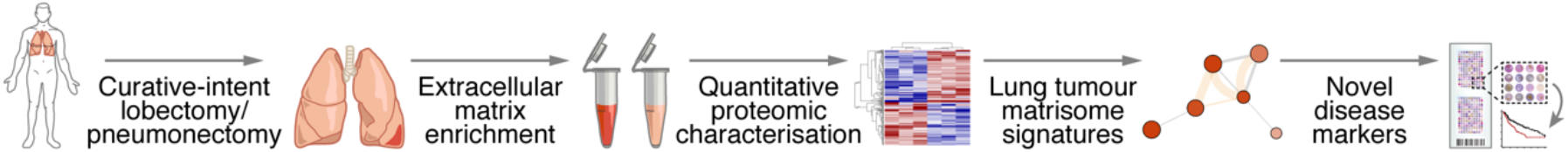

## Introduction

Lung cancer, including non-small cell lung cancer (NSCLC), is a common cancer and the leading cause of cancer-related deaths (Bray et al., 2018). Lung cancer, therefore, is a significant cause of morbidity and mortality, and improving lung cancer outcomes remains a clinically unmet need. Underpinning improved outcomes are a better understanding of the underlying biological process of tumour progression, identification of novel drug targets and establishment of markers that aid clinicians in determining diagnosis, prognosis and treatment decisions. Thus, proteomic assessment of primary cancer samples may help to advance these aims by uncovering differences in the protein composition of tumour and non-tumour tissues (Indovina et al., 2013; Simpson et al., 2008).

The extracellular matrix (ECM) proteome, or matrisome, is composed of core ECM proteins, including collagens, glycoproteins and proteoglycans which form the structure of the ECM, and ECM-associated proteins, such as mucins, enzymes that can modify the ECM and secreted factors such as cytokines (Hynes and Naba, 2012). Matrisome proteins have several key biological functions, including providing physical support, regulating pH, hydration and organisation of tissue and modulating signalling in tissue by binding growth factors and cytokines (Frantz et al., 2010; Hynes, 2009). The matrisome is frequently altered in neoplastic tissues, impacting on multiple hallmark features of cancer (Hanahan and Weinberg, 2011, Pickup et al., 2014); for example, ECM dysregulation is associated with suppression of anti-tumour immunity and immunotherapy resistance, including in lung cancer (Chakravarthy et al., 2018; Peng et al., 2020). Despite having important functions in cancer progression, the matrisome is an under-explored region of tumour tissues (Byron et al., 2013; Filipe et al., 2018; Taha and Naba, 2019), owing in part to technical difficulties in its analysis, including the poor solubility of many ECM proteins (Randles et al., 2017), the multiplicity of their post-translational modifications (Dengjel et al., 2020), the limited number of robustly validated antibodies targeting ECM proteins (Rickelt and Hynes, 2018) and low abundance of ECM proteins in many tissues compared to intracellular proteins (Lindsey et al., 2018). While bulk lung tumour tissue samples have been analysed by mass spectrometry (MS)-based proteomics to search for candidate biomarkers of disease (Hsu et al., 2016; Kikuchi et al., 2012; Nigro et al., 2015; Sandri et al., 2018; Zeng et al., 2012; Zhou et al., 2021), the tumour-associated matrisome of lung cancer patients has not been comprehensively documented. We hypothesised that deep, quantitative characterisation of ECM isolated from tumours from NSCLC patients will enable the detection of new prospective extracellular protein markers of lung cancer.

This study aimed to detect changes in matrisome proteins between human NSCLC tissues and patient-matched non-cancerous lung using a quantitative proteomic approach to curate an extensive resource of lung tumour matrisome proteins. We analysed tissue from 34 patients undergoing curative intent resections for NSCLC, coupling MS-based proteomics to fractionation of tissue samples to enrich for matrisome proteins. We quantified the differential abundance of proteins between tumour tissue and non-cancerous lung, characterising tumour matrisome signatures in NSCLC. Functional network analysis identified a collagen hydroxylation module enriched in tumour tissue, and we validated two novel putative extracellular markers, peroxidasin and a disintegrin and metalloproteinase with thrombospondin motifs 16 (ADAMTS16), as being up-regulated within tumours in separate patient cohorts.

## Results

### NSCLC proteomic study population characteristics

The cohort comprised 34 NSCLC patients, 20 female and 14 male, ranging from 49 to 87 years old, whom underwent surgical resections of NSCLC as treatment with curative intent (details in Supplementary Table 1). Seventeen of the patients were diagnosed with adenocarcinomas, 12 with squamous cell carcinomas, three with large cell tumours and two with pleomorphic tumours. Using TNM classification (8^th^ edition), one patient had a T1, 17 had T2, 10 had T3 and six had T4 tumours (Detterbeck et al., 2017). Lymph node (N) component included stage N0 in 24 patients, seven patients with N1 and three with N2 disease. As this was a curative intent cohort, no patients had distant metastasis. All patients except one were either previous or current cigarette smokers. The standardised uptake value of the PET imaging tracer 18F-FDG from the pre-operative CT scans of patients was high for all but two patients (one moderate, one not recorded). Non-cancerous lung tissue was retrieved from adjacent areas of resected lung tissue removed with the lung tumours (Fig. 1a).

**Fig. 1.**
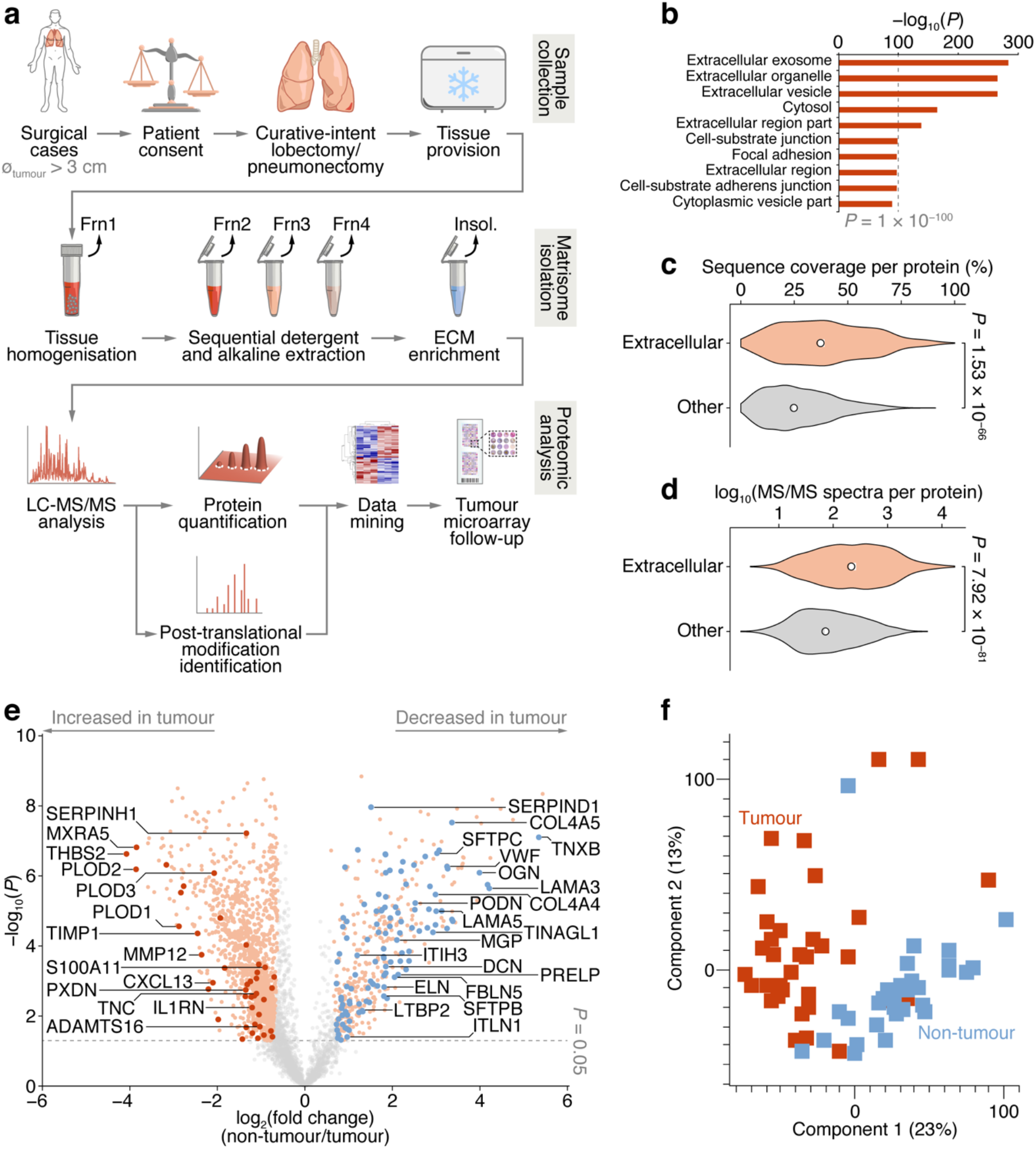
Proteomic analysis of patient-derived lung tumour ECM. **a** Summary of the workflow for the collection of tissue samples, isolation of insoluble (insol.) ECM proteins and MS-based proteomic analysis of the lung tumour matrisome. Frn, fraction. **b** Over-representation analysis of GO cellular component terms in the dataset of proteins quantified by MS-based proteomics. The top ten most enriched terms are shown (*P* < 0.01, Fisher’s exact test with Benjamini–Hochberg correction). All enriched terms are provided in Supplementary Data 2. **c** Distribution of proportions of protein sequences (proportions of possible tryptic peptides per protein) covered by unique peptides identified by MS. **d** Distribution of log_10_-transformed spectral counts per protein. Statistical analysis for **c** and **d**, two-sided Welch’s *t*-test (*n* = 1,420 and 2,182 proteins for extracellular proteins (those annotated with GO term extracellular region) and other proteins, respectively). Black circle, median; white bar, 95% confidence interval; silhouette, probability density. **e** Volcano plot of lung tissue proteins quantified by MS-based proteomics. Significantly differentially regulated proteins enriched or depleted by at least two-fold are coloured orange; differentially regulated matrisome proteins are indicated with large coloured circles (red, increased in tumour; blue, decreased in tumour) (*P* < 0.05, paired two-sided Student’s *t*-test with Benjamini–Hochberg correction). Selected proteins are labelled with gene names for clarity. **f** Principal component analysis of lung tissue samples analysed by MS-based proteomics.

Histopathological abnormalities within the non-malignant regions of the lung included single abnormalities or combinations of pneumonia, emphysema, inflammation, sarcoidosis and pleural fibrosis, and seven of the tissues were recorded as being histopathologically normal lung (Supplementary Table 1).

### Quantification of NSCLC matrisome proteins by MS

To enrich extracellular proteins from patient-derived lung tissue for proteomic analysis, we used detergent and alkaline extractions and DNase treatment to deplete the lung tissue of cells and intracellular material, with minimal loss of relatively insoluble ECM proteins (Fig. 1a and Supplementary Fig. 1a). We performed label-free liquid chromatography-coupled tandem mass spectrometry (LC-MS/MS) analysis of ECM isolated from NSCLC tumour and non-tumour tissue, identifying a total of 1,662,859 peptide–spectrum matches, quantifying 3,602 proteins with a false discovery rate (FDR) of 1% in at least 33% of samples (Supplementary Data 1). Extracellular proteins were significantly over-represented in the isolated ECM fractions (Fig. 1b and Supplementary Data 2), and 1,420 identified proteins (39.4%) were annotated as extracellular in the Gene Ontology database (The Gene Ontology Consortium, 2019) (Supplementary Data 1, 2). The median sequence coverage of extracellular proteins, determined by MS, was significantly higher than that of all other identified proteins (Fig. 1c), and the spectral counts for extracellular proteins were significantly higher than for other proteins (Fig. 1d).

Two hundred and fifty-six proteins in the dataset were classified as curated matrisome proteins (Naba et al., 2016, Shao et al., 2019), which comprised 108 core matrisome proteins and 148 matrisome-associated proteins. The core matrisome group consisted of 70 glycoproteins, 26 collagens and 12 proteoglycans; the matrisome-associated group consisted of 37 ECM-affiliated proteins, 73 ECM regulators and 38 secreted proteins. This is comparable in number and matrisome subgroup distribution to recent proteomic analyses of lung tumour tissue (Gocheva et al., 2017; Tenzer et al., 2016; Tian et al., 2019) (Supplementary Fig. 1b and Supplementary Table 2). Together, these data indicate strong detection and enrichment of extracellular proteins in the patient-derived lung ECM samples.

Of the 3,602 identified proteins, 1,805 significantly differed by at least 2-fold between tumour and non-tumour tissues (*P* < 0.05, paired two-sided Student’s *t*-test with Benjamini–Hochberg correction) (Fig. 1e and Supplementary Data 3). Pairwise correlation analysis of all tumour and non-tumour samples revealed positive correlation between like sample types (Supplementary Fig. 1c), and tumour samples generally clustered together (Fig. 1f and Supplementary Fig. 1c). One hundred and sixty-one matrisome proteins were significantly differentially expressed (Supplementary Data 4), 47 of which were increased in tumour samples compared to non-tumour samples, and 114 of which were decreased in tumour samples compared to non-tumour samples (Fig. 2 and Supplementary Fig. 2).

**Fig. 2.**
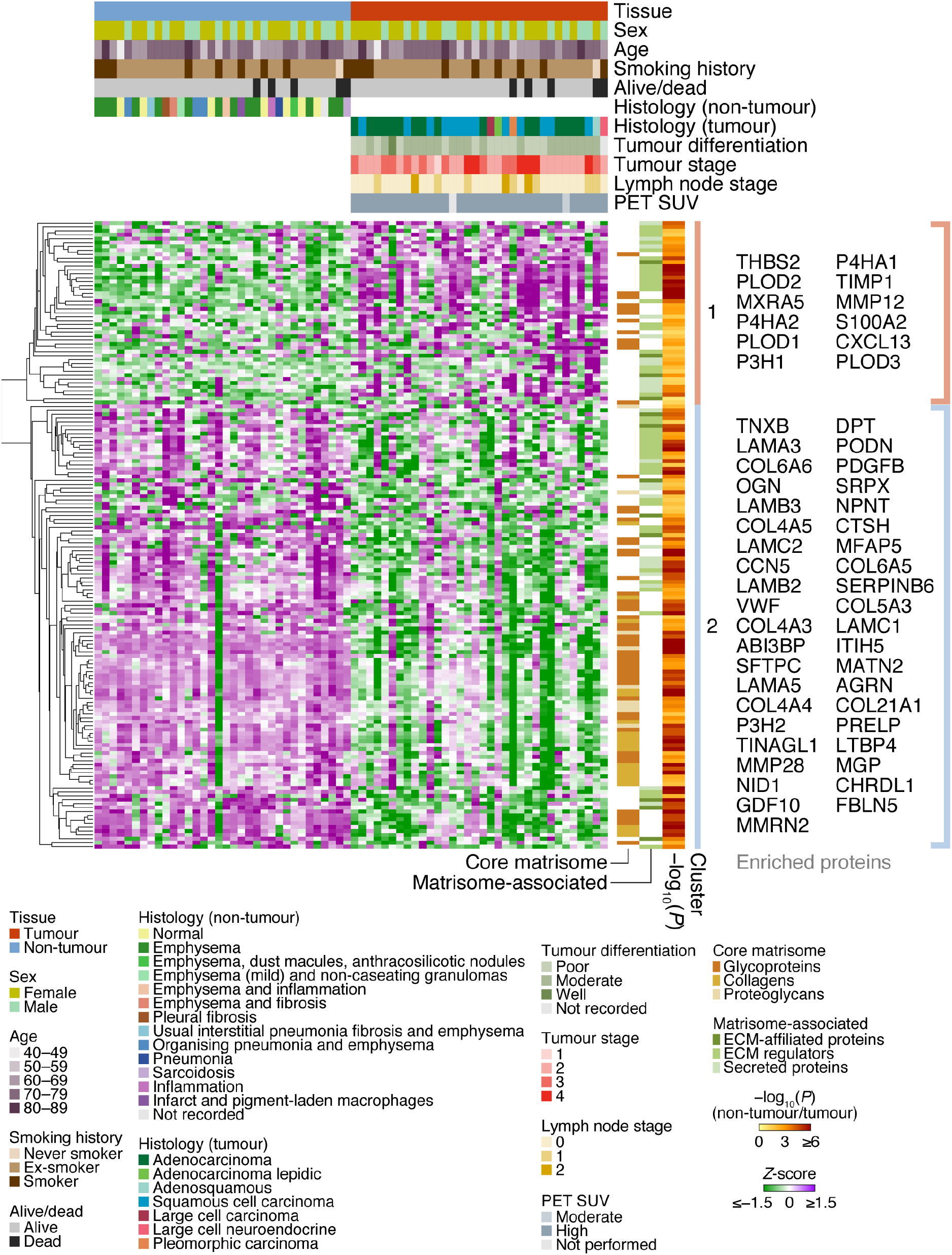
Cluster analysis of patient characteristics and matrisome protein expression. Heatmap representation of the 161 differentially expressed matrisome proteins (*P* < 0.05, paired two-sided Student’s *t*-test with Benjamini–Hochberg correction). Proteins were quantified by log_2_-transformed label-free quantification intensities, standardised by row-wise *Z*-scoring and hierarchically clustered on the basis of Euclidean distance. The two principal clusters are indicated; proteins enriched or depleted by at least four-fold are labelled next to the respective cluster (selected proteins are labelled with gene names for clarity). SUV, standardised uptake value.

### Down-regulation of ECM-organising proteins in NSCLC

The majority of matrisome proteins up-regulated in tumour ECM were matrisome-associated proteins, predominantly ECM regulators and secreted proteins, whereas proteins down-regulated in tumour ECM included a substantial number of both matrisome-associated proteins and core matrisome proteins (Fig. 2 and Supplementary Fig. 2). Of those collagens that were differentially regulated, all were down-regulated in patient-derived tumour samples compared to matched non-tumour samples, ranging from 1.7-fold for type I collagen α1 chain (COL1A1) to 18.1-fold for type VI collagen α6 chain (COL6A6) (Supplementary Fig. 2 and Supplementary Data 3). In addition, all detected laminins were down-regulated in tumour samples compared to matched non-tumour samples, from 1.8-fold for laminin α2 chain (LAMA2) to 18.6-fold for laminin α3 chain (LAMA3) (Supplementary Fig. 2 and Supplementary Data 3). These findings agree with similar observations of decreased abundance of many collagen and laminin subunits in a murine model of lung adenocarcinoma (Gocheva et al., 2017).

Matrisome proteins down-regulated in tumours were enriched for ECM organisation functions (Supplementary Data 5), and these proteins formed a functional subnetwork dominated by collagens and laminins that clustered based on protein interactions (Fig. 3a and Supplementary Fig. 3a). Other down-regulated glycoproteins involved in ECM organisation clustered together, including elastin (3.5-fold), EMILIN-1 (2.0-fold), fibulin-5 (4.1-fold) and microfibril-associated glycoprotein 4 (2.8-fold) (Fig. 3a and Supplementary Fig. 3a), all of which have been previously reported to be down-regulated in lung tumour ECM compared to normal ECM in mice (Gocheva et al., 2017). These data suggest that core matrisome proteins with key roles in ECM organisation and structural integrity are dysregulated in NSCLC tumours.

**Fig. 3.**
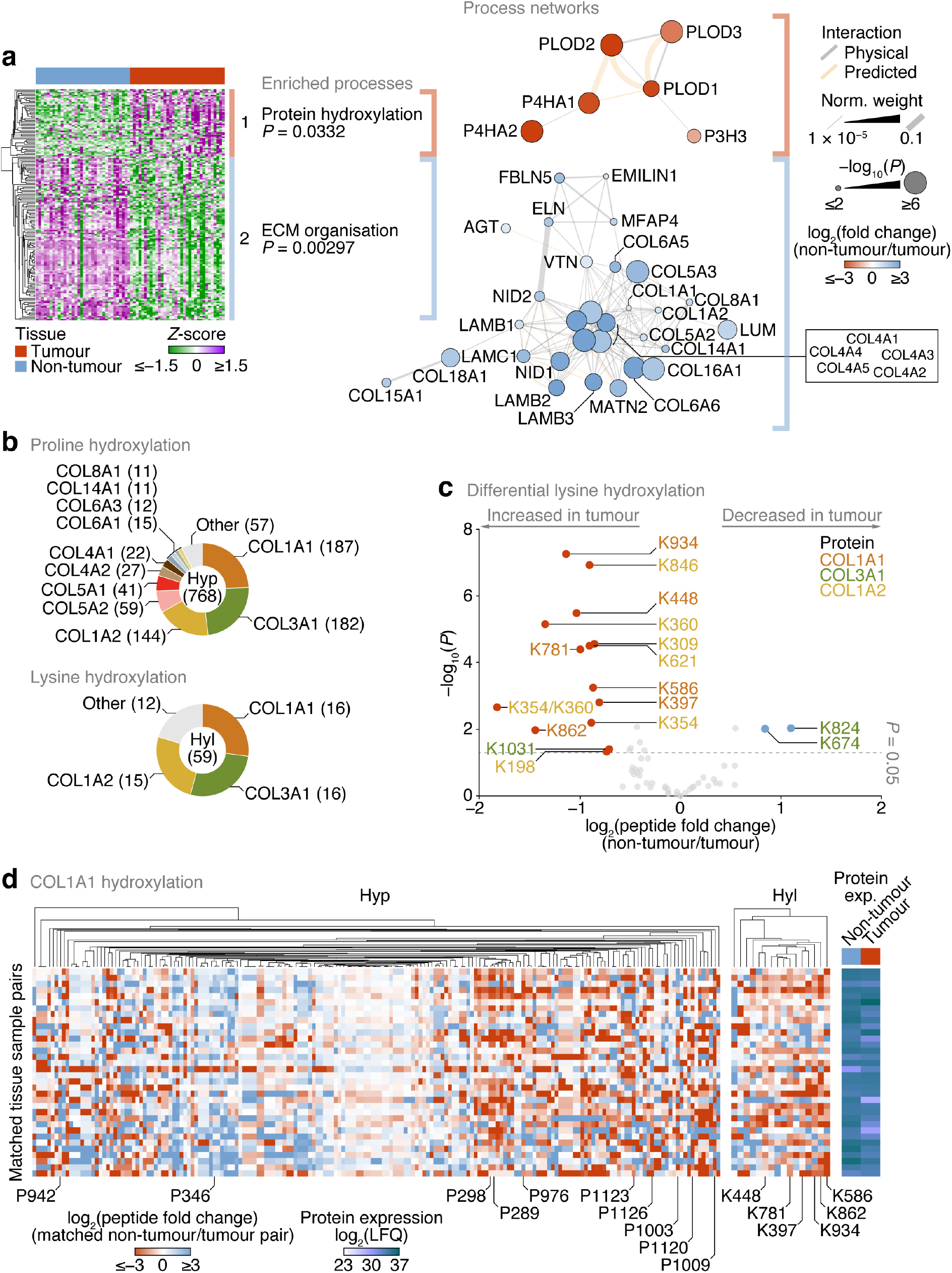
Up-regulation of lysine hydroxylation in patient-derived lung tumour ECM. **a** Gene ontology enrichment analysis of the principal clusters of differentially regulated matrisome proteins identified in Fig. 2. All enriched terms for each respective cluster are shown (Fisher’s exact test with Benjamini–Hochberg correction); for cluster 2, the term “extracellular structure organisation”, the parent term of ECM organisation, and comprising the same contributing proteins and enriched with the same *P*-value, was omitted. Proteins belonging to enriched terms were used to construct corresponding protein interaction networks (right panels). Proteins (nodes) are coloured according to enrichment or depletion in tumour samples and sized according to statistical significance (*P* < 0.05, paired two-sided Student’s *t*-test with Benjamini–Hochberg correction). Protein interactions (edges) were weighted according to evidence of co-functionality. Unconnected proteins are not shown. **b** Proportions of identified peptides containing hydroxylated proline (Hyp; top panel) or hydroxylated lysine (Hyl; bottom panel) assigned to corresponding proteins. Numbers of modified peptides are indicated in parentheses. **c** Volcano plot of peptides containing hydroxylated lysine quantified by MS-based proteomics. Differentially regulated peptides are indicated with large coloured circles (red, increased in tumour; blue, decreased in tumour) (*P* < 0.05, FDR 20%, paired two-sided Student’s *t*-test with Benjamini–Hochberg correction). **d** Regulation of proline and lysine hydroxylation in type I collagen α1 chain (COL1A1) across 34 matched non-tumour–tumour paired tissue samples. The top ten most enriched peptides for each hydroxylation modification are labelled (all six enriched peptides containing hydroxylated lysine are labelled). Total protein expression determined by label-free quantification (LFQ) shown for corresponding samples.

### Increased lysine hydroxylation in fibrillar collagens in NSCLC

Matrisome proteins up-regulated in tumours were enriched for protein hydroxylation processes (Supplementary Data 5), driven by procollagen-lysine,2-oxoglutarate 5-dioxygenases (also known as lysyl hydroxylases) and prolyl 3-and prolyl 4-hydroxylases (Supplementary Fig. 2 and Supplementary Data 3), which formed a functional subnetwork of interacting proteins (Fig. 3a and Supplementary Fig. 3a). These proteins catalyse the post-translational formation of hydroxylysine and hydroxyproline residues in collagen chains, which are critical for collagen helix stability (Bella and Hulmes, 2017), and have been reported to be up-regulated in fibrotic lung ECM in murine and human model systems (Schiller et al., 2015; Merl-Pham et al., 2019). LC-MS/MS analysis revealed that the majority of hydroxylated residues were found in type I and type III collagen chains (COL1A1, COL1A2, COL3A1) (Fig. 3b). While there were both up- and down-regulated hydroxyproline-containing peptides quantified by LC-MS/MS (Supplementary Fig. 3b), almost all hydroxylysine-containing peptides were up-regulated in tumour ECM, and all-but-one of these were derived from type I collagen chains (Fig. 3c).

To identify clusters of residues that showed similar regulation of hydroxylation across the tissue samples, we mapped the changes in abundance of hydroxyproline- and hydroxylysine-containing peptides for matched tumour–non-tumour pairs. This analysis indicated variable modulation of proline hydroxylation in type I and type III collagen chains, including a substantial cluster of hydroxyproline-containing peptides derived from COL1A1 that were up-regulated in tumour ECM (Fig. 3d and Supplementary Fig. 3c, d). Lysine hydroxylation, however, was predominantly up-regulated in type I collagen chains across most matched tumour samples (Fig. 3c, d and Supplementary Fig. 3d), whereas modulation of lysine hydroxylation in type III collagen was more variable (Fig. 3c and Supplementary Fig. 3c). These data imply that type I collagen lysine residues are extensively hydroxylated in tumour ECM, in concert with an up-regulation of enzymes that catalyse collagen hydroxylation.

### Up-regulation of matrisome-associated proteins in NSCLC

In addition to enrichment of enzymes that regulate ECM hydroxylation in tumours tissue, there was an increase in matrisome proteins associated with ECM turnover in tumour samples. Several cathepsins, some of which have reported roles in the degradation of extracellular proteins, were up-regulated in tumour samples, including the secreted thiol proteases cathepsin B (2.1-fold) and cathepsin S (1.7-fold) (Supplementary Fig. 2 and Supplementary Data 3). Several matrix metalloproteinases (MMPs) were up-regulated in tumour samples, namely MMP12 (5.1-fold), MMP14 (2.3-fold) and MMP2 (2.2-fold), which cleave elastin (and insulin), collagen (and other extracellular proteins, such as aggrecan) and certain collagens (and gelatin), respectively. In contrast MMP28, which cleaves casein, was down-regulated in tumour samples (7.3-fold) (Supplementary Fig. 2 and Supplementary Data 3). A disintegrin and metalloproteinase with thrombospondin motifs 16 (ADAMTS16), another protease, was up-regulated in tumour samples (2.0-fold).

In addition to many matrisome-associated proteins, several core matrisome glycoproteins were up-regulated in tumour tissue. The most enriched glycoproteins in tumour ECM compared to non-tumour ECM included thrombospondin-2 (16.9-fold), MXRA5 (14.4-fold), peroxidasin (2.5-fold), thrombospondin-1 (2.5-fold), SPARC (2.3-fold), FRAS1 (2.3-fold) and tenascin-C (2.1-fold) (Supplementary Fig. 2 and Supplementary Data 3). Four of the top five most up-regulated glycoproteins in tumour ECM – thrombospondin-1 and -2, SPARC and peroxidasin – formed a connected subnetwork (Supplementary Fig. 3a), pointing to their common functional roles in organising the collagen ECM and regulating cell–ECM adhesion.

### Peroxidasin and ADAMTS16 are increased in NSCLC tumour samples

Of the extracellular proteins we quantified as up-regulated in tumours, we selected for further analysis two – peroxidasin and ADAMTS16 (Fig. 4a) – that are not well understood in NSCLC. We constructed a tumour microarray (TMA) containing control and tumour samples from a separate cohort of lung cancer patients undergoing curative intent surgery (Supplementary Table 3). Selected proteins were analysed by immunohistochemistry (IHC) and scored on a four-point scale (scored from 0–3) based on increasing immunostaining by IHC from no staining to strong staining. For lung cores in the TMA derived from tumour samples, the distribution and scoring of immunostained proteins was assessed separately for tumour stroma and tumour cells. In tumours, peroxidasin and ADAMTS16 were found variably distributed between tumour stroma and tumour cells (Fig. 4b).

**Fig. 4.**
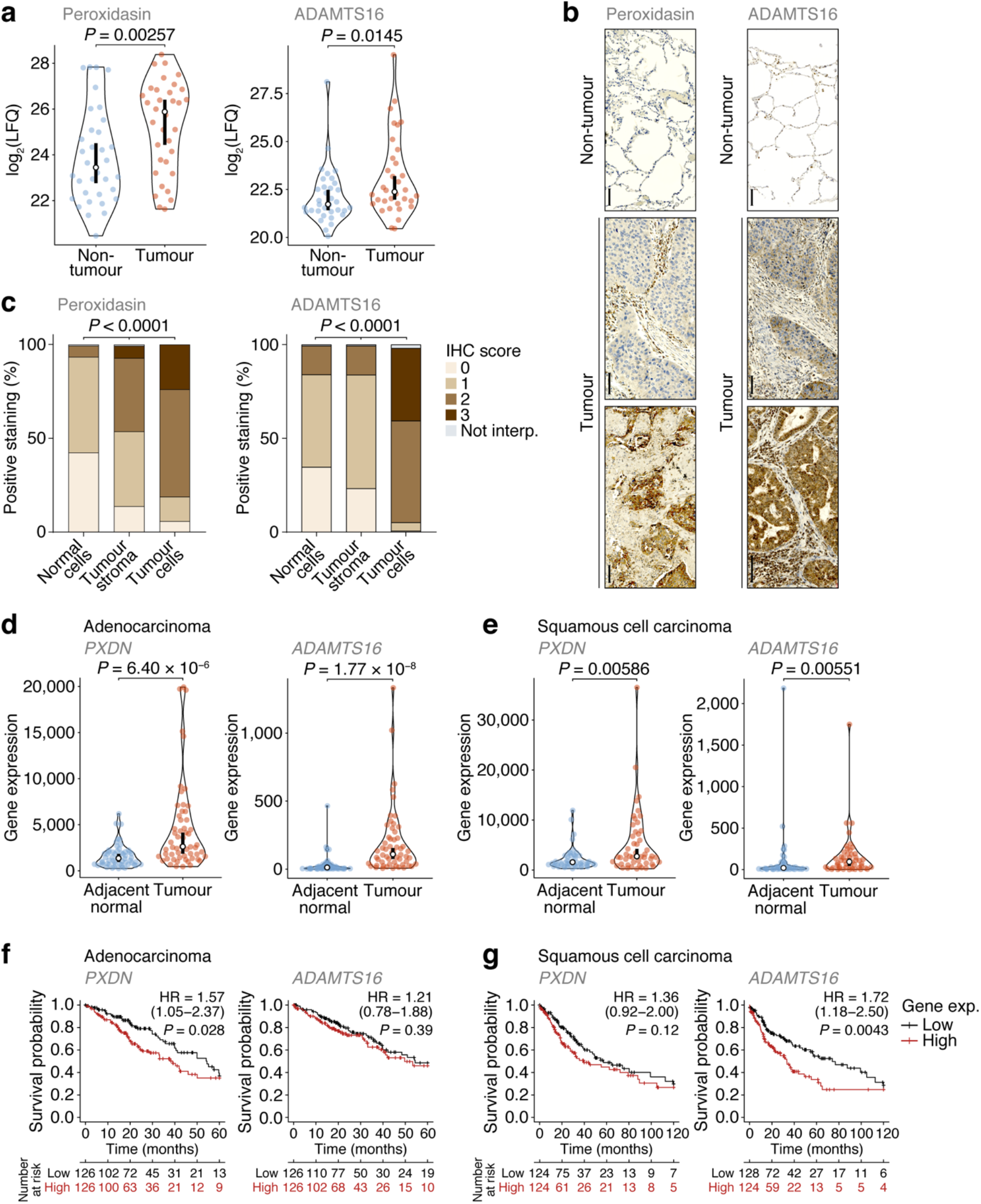
Dysregulation of peroxidasin and ADAMTS16 in NSCLC tumour cells. **a** Label-free quantification of peroxidasin (left panel) and ADAMTS16 (right panel) in non-tumour and tumour ECM samples quantified by MS-based proteomics. Statistical analysis, paired two-sided Student’s *t*-test with Benjamini–Hochberg correction (*n* = 34 matched non-tumour–tumour paired tissue samples). **b** Immunohistochemistry (IHC) anaysis of peroxidasin (left panels) and ADAMTS16 (right panels) for exemplar lung cores from the tumour microarray. Scale bars, 100 μm. **c** Quantification of TMA immunostaining for non-tumour-derived lung cores (normal cells) and tumour-derived lung cores (tumour stroma and tumour cells). IHC scores of 0, 1, 2 and 3 correspond to no, mild, moderate and strong immunostaining, respectively. Not interp., not interpretable. Statistical analysis, chi-square test. **d, e** Differential expression analysis of *PXDN* (left panels) and *ADAMTS16* (right panels) in cohorts of lung adenocarcinoma (**d**) or lung squamous cell carcinoma (**e**) patients from integrated RNA-seq datasets. Statistical analysis, Wilcoxon signed-rank test (*n* = 57 and 49 matched adjacent normal–tumour paired tissue samples for adenocarcinoma and squamous cell carcinoma patients, respectively). For **a, d** and **e**, black circle, median; black bar, 95% confidence interval; silhouette, probability density. **f, g** Kaplan–Meier curves of patient survival associated with degree of expression of *PXDN* (left panels) and *ADAMTS16* (right panels) in cohorts of lung adenocarcinoma (**f**) or lung squamous cell carcinoma (**g**) patients from TCGA RNA-seq datasets. The lower quartiles (low) and upper quartiles (high) of gene expression were compared. HR, hazard ratio (with 95% confidence interval). Statistical analysis, log-rank test (*n* = 513 and 501 patients for adenocarcinoma and squamous cell carcinoma cohorts, respectively).

Importantly, both proteins had significantly increased immunostaining intensities in tumour cells compared to non-tumour cells (Fig. 4b, c). These data confirm the results from the MS-based proteomics analysis and imply that peroxidasin and ADAMTS16 are up-regulated in NSCLC tumours.

To examine the expression of the genes encoding the selected extracellular proteins, we analysed an integrated dataset of RNA-seq data derived from paired tumour and adjacent normal tissue from lung adenocarcinoma or lung squamous cell carcinoma patients (Bartha and Győrffy, 2021). Expression of both *PXDN* (which encodes peroxidasin) and *ADAMTS16* were significantly up-regulated in adenocarcinoma tumours compared to matched adjacent normal lung tissue (1.96-fold and 10.7-fold change in median expression, respectively) (Fig. 4d). Both genes were also significantly up-regulated in squamous cell carcinoma tumours compared to matched adjacent normal lung tissue (1.78-fold and 5.62-fold change in median expression, respectively) (Fig. 4e). These data indicate that *PXDN* and *ADAMTS16* are transcriptionally up-regulated in NSCLC tumours.

We next assessed whether the protein expression levels of peroxidasin and ADAMTS16 were predictors of survival in the TMA cohort (Supplementary Table 3). Although up-regulated in tumour tissue (Fig. 4a–c), the degree of tumour stroma immunostaining of these two markers, as determined by IHC scoring of all interpretable lung cores, did not significantly correlate with survival outcome within the limited TMA cohort (Supplementary Fig. 4). To examine patient survival in larger NSCLC cohorts, we used the Cancer Genome Atlas (TCGA) data derived from primary tissue samples from lung adenocarcinoma (513 patients) or lung squamous cell carcinoma (501 patients). We found that higher expression of *PXDN* was significantly associated with shorter survival of adenocarcinoma patients (hazard ratio (HR) = 1.57; 95% confidence interval (CI), 1.05–2.37; *P* = 0.028, log-rank test), whereas that of *ADAMTS16* was not (Fig. 4f). Median survival of adenocarcinoma patients was 54.4 months for the low *PXDN* expression subgroup and 39.0 months for the high *PXDN* expression subgroup. In contrast, there was no statistical association of expression of *PXDN* with survival of squamous cell carcinoma patients, whereas higher expression of *ADAMTS16* was significantly associated with shorter survival of squamous cell carcinoma patients (HR = 1.72; 95% CI, 1.18–2.50; *P* = 0.0043) (Fig. 4g). Median survival of squamous cell carcinoma patients was 74.1 months for the low

*ADAMTS16* expression subgroup and 33.4 months for the high *ADAMTS16* expression subgroup. Together, these results indicate that these proteins, identified in the patient lung tumour matrisome, are increased in NSCLC and their corresponding gene expression provides prognostic information for lung adenocarcinoma (*PXDN*) or squamous cell carcinoma (*ADAMTS16*) patients undergoing curative intent surgery.

## Discussion

We present herein a comprehensive characterisation of the NSCLC matrisome. To our knowledge, this is the first unbiased matrisome-scale proteomic analysis of enriched ECM from lung cancer patient tissue. Our quantitative analyses reveal the differential expression of a substantial number of extracellular proteins in lung tumour tissue from patients with early-stage NSCLC, implying extensive remodelling of the extracellular niche.

We found that many core matrisome proteins were less abundant in lung tumour tissue as compared to non-cancerous lung tissue. For example, all differentially expressed collagens and proteoglycans were relatively decreased in tumour samples, while a subset were not significantly altered. These findings are consistent with MS-based proteomic data from a murine model of lung adenocarcinoma, which identified decreased or little change in abundance of many core matrisome proteins, including the majority of detected collagens and laminins, in tumour ECM as compared to non-tumour ECM (Gocheva et al., 2017).

The general association of lung cancer with desmoplasia, however, suggests that our observations may be linked to early stages of tumour ECM remodelling prior to accumulation of collagen. Indeed, recent examination of idiopathic pulmonary fibrosis lung tissue identified induction of collagen-modifying enzymes that contribute to collagen cross-linking, including lysyl hydroxylase 2, but not increased collagen synthesis, as a defining feature of lung fibrosis that increases tissue stiffness and promotes fibrotic progression (Brereton et al., 2022; Jones et al., 2018). Lysyl hydroxylase 2 (encoded by *PLOD2*) is one of several collagen-modifying enzymes that we detected as up-regulated in NSCLC tumour tissue in this study, alongside a concomitant increase in collagen lysine hydroxylation. Lysyl hydroxylase 2 is secreted by lung cancer cells in culture, and its hydroxylation of collagen telopeptidyl lysine residues leads to the formation of stable hydroxylysine aldehyde-derived collagen cross-links that are up-regulated in lung cancer tissue and generate stiffer tumour tissue (Chen et al., 2015; Chen et al., 2016). Thus, dysregulation of collagen architecture or cross-linking, and consequential ECM stiffness, may be more prominent features of early-stage NSCLC than changes in total collagen synthesis or density.

We observed the up-regulation of enzymes involved in extracellular protein degradation in lung tumour tissue, including several cathepsins and MMPs. These proteinases target a wide range of ECM proteins, such as collagen, laminin and elastin (Lu et al. 2011; Vizovišek et al., 2019), almost all of which were depleted in tumour samples. This suggests that there may be increased turnover of core ECM macromolecules in NSCLC tumours, consistent with the remodelling of the extracellular niche found in various respiratory diseases and invasive tumour growth (Burgess et al., 2016; Lu et al., 2011). Indeed, the expression of MMP2 and MMP14, which were up-regulated in lung tumour ECM, is linked to poorer patient outcomes in NSCLC (Passlick et al., 2000; Wang et al., 2014), and MMP12, also up-regulated in tumour samples, is associated with faster disease relapse and metastasis in NSCLC patients (Hofmann et al., 2005) and the occurrence of bronchioalevolar adenocarcinomas in patients with emphysema (Qu et al., 2009).

We validated in a separate patient cohort the up-regulation in tumour tissue of two extracellular proteins identified by MS-based proteomics. Peroxidasin, an extracellular peroxidase, mediates the formation of sulfilimine cross-links between methionine and hydroxylysine residues in type IV collagen (Bhave et al., 2012; Lázár et al., 2015). Interestingly, the activity of peroxidasin is associated with accelerated tumorigenesis in murine models of lung carcinoma in the presence of inhalable fine particulate matter, which adsorbs peroxidasin and leads to accumulation of collagen cross-linking (Wang et al., 2022). In our analyses, peroxidasin clustered in a tumour-enriched collagen-modifying protein subnetwork, which together with our identification of a tumour-enriched collagen hydroxylation functional network, implies that collagen modifications and modulation of collagen cross-linking are key characteristics of early-stage NSCLC. The other selected extracellular tumour marker candidate, ADAMTS16, a matrisome-associated protease, targets fibronectin to inhibit ECM assembly (Schnellmann et al., 2018), further suggesting that ECM organisation is dysregulated in NSCLC tissue. IHC analyses determined that both peroxidasin and ADAMTS16 were enriched in tumour cell-rich regions of lung tumour tissue, although IHC scoring of either of these candidates did not represent a significant predictor of survival in the TMA cohort of NSCLC patients. The assessment of patients with early-stage disease undergoing curative intent surgery in this study precludes the detection of matrisome protein changes that occur in more advanced disease, which could limit the ability to identify robust late-stage disease biomarkers but likely also explains the identification of putative early events in the remodelling of collagen cross-linking. In addition, tissue was only collected from a small portion of the tumour. Tumours are known to be heterogeneous (Bedard et al., 2013; Marusyk et al., 2012); therefore, sampling from one area may not be representative of all the pathological protein changes within a tumour. Analysis of transcriptomic data from larger lung cancer patient cohorts, however, revealed increased tumour expression of genes encoding both peroxidasin and ADAMTS16, and this was separately linked to poorer survival of lung adenocarcinoma and squamous cell carcinoma patients, respectively. Recent analysis of matrisome gene expression in multiple transcriptomic datasets showed that extracellular protein levels are generally concordant with corresponding gene expression in human tissues (Nieuwenhuis et al., 2021). Our findings are consistent with this observation and suggest that high expression of *PXDN* and *ADAMTS16* can be used as surrogate readouts for up-regulation of these two potential extracellular lung tumour markers.

In summary, this study provides an extensive analysis of lung tissue ECM in patients with lung cancer, charting the remodelling of the matrisome in early-stage NSCLC. We show that proteomic profiling of patient-derived lung tumour ECM enables the identification of candidate extracellular markers of tumour cells. In addition, the systems-level changes to the lung matrisome we report here, including the up-regulation of a collagen hydroxylation network in NSCLC tissue, reveal potential molecular networks that could modulate ECM organisation and regulate lung cancer progression.

## Materials and Methods

### Study approvals

Patient samples were collected with ethical approval and written patient consent. Ethical approval was granted by Lothian NRS Bioresource, REC number 15/ES/0094 (reference SR419). All samples were assigned an anonymised code, and researchers were blinded to patient details for experiments. The TMA dataset was approved by Lothian NRS Bioresource, REC number 15/ES/0094 (reference SR1208), and approved by the NHS Lothian Caldicott Guardian (reference CRD19031).

### Patient samples

Tissues samples used for mass spectrometry (MS) were collected from patients with NSCLC undergoing curative-intent surgical procedures. Following resection, samples were handled by an experienced thoracic pathologist, and samples of tumour and non-cancerous lung (from the most distal portion of the resection specimen) were dissected and provided for MS analysis. Samples were snap frozen and stored at −70°C until required. Anonymised patient details were recorded, including age, gender, smoking history, histopathological diagnosis of tumour and non-tumour tissues, degree of differentiation of the tumour tissue, tumour stage and lymph node stage (determined by TNM status), survival and PET tracer uptake.

### Enrichment of matrisome proteins

Tissue samples were enriched for predominately insoluble matrisome proteins by depleting soluble intracellular proteins (Fig. 1a). Methods were adapted from previously published work (Lennon et al., 2014). Tissue samples were finely minced with scalpels, and the presence of any necrosis and tissue pigmentation was noted. Samples were homogenised twice in 1 ml of chilled phosphate-buffered saline (PBS) (without Ca^2+^ or Mg^2+^), containing 1% (v/v) protease inhibitor cocktail (Sigma-Aldrich), using a Precellys 24 tissue homogeniser (Bertin Instruments) at 6,500 rpm for 50 s. Samples were incubated for 5 min on ice between homogenisation cycles. The homogenate was centrifuged at 14,000 × *g* for 10 min at 4°C. The supernatant was removed (fraction 1), and the remaining pellet was resuspended and incubated in 10 mM Tris-HCl, pH 8, 150 mM NaCl, 25 mM EDTA, 1% (v/v) Triton X-100, 1% (v/v) protease inhibitor cocktail for 30 min on ice. The lysate was centrifuged at 14,000 × *g* for 10 min at 4°C. The supernatant was removed (fraction 2), and the remaining pellet was resuspended and incubated in 20 mM NH_4_OH containing 0.5% (v/v) Triton X-100 in PBS (without Ca^2+^ or Mg^2+^). The lysate was centrifuged at 14,000 × *g* for 10 min at 4°C. The supernatant was removed (fraction 3), and the remaining pellet was incubated with 10 μg/ml DNase I in PBS (with Ca^2+^ and Mg^2+^) for 30 min on ice. Samples were centrifuged at 14,000 × *g* for 10 min at 4°C, and the supernatant was removed (fraction 4). The remaining pellet was washed in ice-cold PBS and centrifuged at 14,000 × *g* for 10 min at 4°C three times. The final insoluble pellets enriched for ECM proteins were then stored at −70°C until further use.

### SDS-PAGE and western blotting

To confirm matrisome proteins were enriched in the final protein pellet prior to proteomics experiments, the supernatants from serially extracted fractions from a subset of samples were probed by SDS-PAGE and western blotting. Three percent of the supernatant from each fraction was used for western blotting to allow comparison between fractions. Fractions 1–4 were incubated in 1× Laemmli buffer containing 50 mM dithiothereitol for 10 min at 95°C. The final ECM pellet was precipitated using TCA–acetone (see below), homogenised and incubated in 8 M urea, 100 mM NH_4_HCO_3_, pH 8, 10 mM dithiothereitol for 30 min at 37°C. Samples were resolved by SDS-PAGE using 4–20% Tris-glycine gels. Proteins were transferred to PVDF membranes using an iBlot 2 dry blotting system (Thermo Fisher Scientific) according to manufacturer’s instructions. Membranes were blocked using milk blocking buffer (5% (w/v) non-fat skimmed milk powder (Marvel) in 1× Tris-buffered saline containing 0.1% (v/v) Tween 20 (TBS-T)) for 1 h at room temperature with shaking. Following blocking, membranes were incubated with primary antibodies, diluted 1:1,000 in milk blocking buffer, overnight at 4°C with rolling. Antibodies used were rabbit polyclonal anti-fibronectin (#ab2413, Abcam), mouse monoclonal anti-lamin A/C (clone 4C11; #4777, Cell Signaling Technology) and rabbit monoclonal anti-GAPDH (clone 14C10; #2118, Cell Signaling Technology). Membranes were washed three times in TBS-T and then incubated with anti-rabbit or anti-mouse horseradish peroxidase-conjugated secondary antibodies, diluted 1:10,000 or 1:5,000, respectively, in milk blocking buffer, for 45 min at room temperature with rolling. Membranes were washed three times in TBS-T. Membranes was developed using Pierce ECL western blotting substrate (Thermo Fisher Scientific) according to the manufacturer’s instructions. Membranes were imaged using an Odyssey Fc imaging system (LI-COR Biosciences).

### Matrisome protein precipitation

Ice-cold TCA (10% (v/v) final concentration) was added to final ECM-enriched fractions for 20 min at 4°C. The sample was centrifuged at 16,000 × *g* for 30 min at 4°C. The supernatant was discarded, and the ECM-enriched pellet was resuspended in ice-cold acetone, using vortexing and sonication, and incubated for 20 min at −20°C. The sample was centrifuged at 16,000 × *g* for 30 min at 4°C, and the supernatant was discarded. The acetone wash step was then repeated. Samples were air dried at room temperature until residual solvent had evaporated. Precipitated protein pellets were stored at −70°C until further use.

### Matrisome protein digestion

Precipitated protein pellets were resuspended in 300 μl solubilisation buffer (8 M urea, 100 mM NH_4_HCO_3_, pH 8, 10 mM dithiothreitol). Samples were sonicated on ice using a probe sonicator and then incubated for 30 min at 37°C. After cooling to room temperature, sample pH in the range pH 8–9 was verified using pH indicator paper. To 50 μl sample, 8.3 μl of 175 mM iodoacetamide in 100 mM NH_4_HCO_3_, pH 8, was added (final concentration 25 mM) and incubated for 30 min at room temperature in the dark to block thiol groups of cysteine residues. Excess iodoacetamide was quenched by the addition of 3 μl of 100 mM dithiothreitol in 100 mM NH_4_HCO_3_, pH 8. For protein digestion, urea was diluted from 8 M to 2 M using 100 mM NH_4_HCO_3_, pH 8. Samples were incubated with 833 units of PNGase F for 2 h at 37°C with shaking (900 rpm), then with 800 ng of Lys-C for 2 h at 37°C, then with 1.6 μg of MS-grade trypsin for 16 h at 37°C with shaking (1,200 rpm). Samples were then incubated with an additional 0.8 μg of trypsin for 2 h at 37°C with shaking (1,200 rpm). Samples were acidified using trifluoroacetic acid (TFA) to obtain sample pH in the range pH 3–4, and samples were clarified by centrifugation at 18,000 × *g* for 15 min.

### Peptide desalting prior to MS analysis

Peptide concentrations in the digested samples were estimated using a Nanodrop spectrophotometer measuring absorbance at 280 nm. Stop-and-go extraction (Stage) tips were made in-house, using a method adapted from Rappsilber et al. (Rappsilber et al., 2003). Briefly, using an 18-gauge blunt-ended needle, two disks were cut from C18 solid-phase extraction material and placed on top of each other inside a 200-μl EasyLoad pipette tip (Greiner Bio-One). Stage tips were loaded into a custom-built tip holder over a deep 96-well plate. Methanol was added to each Stage tip, and tips were centrifuged at 300 × *g* for 2 min. Stage tips were then equilibrated with 0.1% (v/v) TFA and centrifuged at 500 × *g* for 5 min. Samples estimated to contain 10 μg of acidified peptide (based on Nanodrop readings) were then added to Stage tips, and tips were centrifuged at 500 × *g* for 5 min. Stage tips with bound protein were stored at −20°C until elution.

### Desalted peptide elution

Peptides were eluted from C18-containing Stage tips with 40 μl of 80% (v/v) acetonitrile, 0.1% (v/v) TFA, and tips were centrifuged at 200 × *g* for 5 min. Acetonitrile was evaporated using a vacuum centrifuge. Samples were adjusted to 15 μl volume with 0.1% (v/v) TFA, and peptide concentration was re-measured using a Nanodrop spectrophotometer measuring absorbance at 280 nm.

### MS data acquisition

‘Bottom-up’ liquid chromatography-coupled tandem MS (LC-MS/MS) was used to elucidate the structure of isolated peptides and to detect post-translational modifications. LC-MS/MS analysis was carried out using an Orbitrap Fusion Lumos Tribrid mass spectrometer (Thermo Fisher Scientific) coupled to an UltiMate 3000 UHPLC Nano (Thermo Fisher Scientific), Aurora C18 column (IonOpticks), column oven (maintained at 50°C; Sonation) and Proxeon Nanospray ion source (Thermo Fisher Scientific). Peptides were injected onto an Aurora C18 column in buffer A (2% (v/v) acetonitrile, 0.5% (v/v) acetic acid) and eluted with a linear 120-min gradient of 2%–45% (v/v) buffer B (80% (v/v) acetonitrile, 0.5% (v/v) acetic acid). Eluting peptides were ionised in positive-ion mode before data-dependent analysis. A dynamic exclusion window of 30 s was enabled and lockmass was not used.

### MS data analysis

The MS data were normalised and quantified using MaxQuant software (version 1.6.10.43) (Tyanova et al., 2016a). MaxQuant quantifies proteins using a label-free technique, which calculates a normalised peptide abundance from ion signal intensities (Cox et al., 2014). Peptide lists were searched against the human UniProt knowledgebase database (version 2019_09) (UniProt Consortium, 2019) and a common contaminants database using the Andromeda search engine implemented in MaxQuant. Cysteine carbamidomethylation was set as a fixed modification, and methionine oxidation, lysine oxidation, proline oxidation, N-terminal deamidation and protein N-terminal acetylation were set as variable modifications, with up to five modifications per peptide.

Peptide identifications were matched between runs if they eluted within a time window of 0.7 min. Enzyme specificity was C-terminal to arginine and lysine, except when followed by proline. A maximum of two missed cleavages were permitted in the database search; minimum peptide length was seven amino acids. At least two peptide ratios were required for label-free quantification, and large label-free quantification ratios were stabilised. Peptide and protein false-discovery rates (FDRs) were set to 1%, determined by applying the target-decoy search strategy implemented in MaxQuant. Proteins matching to the common contaminants or decoy databases, and matches only identified by site, were omitted. Label-free quantification intensities were log_2_ transformed, and proteins quantified in less than one-third of samples were removed. Missing values were imputed from a width-compressed, down-shifted Gaussian distribution using Perseus (version 1.6.2.3) (Tyanova et al., 2016b).

### Functional enrichment analysis

Gene Ontology (GO) over-representation analysis was performed using DAVID 2021 (DAVID Knowledgebase, version 2022q2) (Sherman et al., 2022). The functional annotation tool category of GO term enrichment was used to filter out very broad GO terms. GO over-representation analysis of matrisome proteins used the entire matrisome database as background. Significant over-representation of terms was determined using a Fisher’s exact test with Benjamini–Hochberg correction.

### Matrisome data analysis

Identified proteins were defined as belonging to the matrisome if they exist in searchable databases of matrisome proteins based on data from 17 studies of the ECM, MatrisomeDB (Shao et al., 2019), and *in-silico* and *in-vivo* data from the Matrisome Project (Naba et al., 2016). Label-free quantification intensities for proteins derived from tumour and non-tumour samples were compared using a paired two-sided Student’s *t*-test, with FDR set to 5% and artificial within-groups variance (s0) set to 1 using Perseus. Statistical data were visualised using Prism (version 9.2.0; GraphPad). For differentially expressed matrisome proteins, potential confounding variables were also considered. These variables included necrotic tumour samples versus non-necrotic tumour samples, adenocarcinoma versus squamous cell carcinoma, well and moderately differentiated tumours versus poorly differentiated tumours, tumours with local lymph node spread versus no known lymph node metastasis and normal non-tumour samples versus non-tumour samples with underlying pathology. Necrosis was crudely assessed based on the composition of the tumour samples when dissected and minced. PET standardised uptake value was scored as mild, moderate or marked. In some instances, absolute values were provided in reports rather than categorical values. These numbers were reclassified as no uptake for 0.6–0.8, low uptake for 1.0–2.0, moderate uptake for 1.5–2.0 and high uptake for >2.5. These variables were compared using multiple Student’s *t*-tests (*P* < 0.05, FDR 5%).

### Hierarchical cluster analysis

To enable relative comparison of protein enrichment, log_2_-transformed label-free quantification intensities were standardised by row-wise (protein-wise) *Z*-scoring. Differentially expressed matrisome proteins were hierarchically clustered on the basis of Euclidean distance, computed using average linkage, using Cluster 3.0 (C Clustering Library, version 1.54) (de Hoon et al., 2004). The following variables were included in the protein enrichment analysis: sex, age, smoking history, if patients had died at the time of analysis, histopathology results, degree of tumour differentiation, tumour stage, lymph node stage and PET 18F-FDG tracer uptake. Modified peptide fold changes between matched non-tumour–tumour pairs were hierarchically clustered on the basis of Euclidean distance, computed using average linkage. For sample correlation analysis, Spearman rank correlation coefficient-based distance matrices were computed using average linkage. Clustering results were visualised using Java TreeView (version 1.1.5r2) (Saldanha, 2004).

### Interaction network analysis

Composite functional association networks were constructed using GeneMANIA (version 3.5.2; human interactions) (Montojo et al., 2010) in Cytoscape (version 3.8.0) (Shannon et al., 2003). Networks were based on reported physical and predicted protein–protein interactions; edges, representing protein–protein interactions, were weighted according to evidence of co-functionality using GeneMANIA. Networks were clustered using Markov clustering (granularity parameter 2.5), and graph layouts were determined using the force-directed algorithm in the Prefuse toolkit (Heer et al., 2005).

### TMA and immunohistochemistry

The TMA was constructed for sequential patients undergoing surgical resection for lung cancer over a 2-year period at a regional thoracic surgery centre. Formalin-fixed paraffin pathological blocks were annotated by an experienced pathologist, and 1-mm cores taken from tumours and non-cancerous lung were embedded into new blocks and, subsequently, 4-μm sections were cut onto glass slides.

Slides were deparaffinised and rehydrated, and antigen retrieval was undertaken with citrate buffer (#ab64214, Abcam) twice for 5 min in a microwave. Slides were processed with a commercial DAB cell and tissue staining kit (#CTS019, R&D Systems). Primary antibody immunostaining was optimised for polyclonal rabbit anti-human peroxidasin (#abx101905, Abbexa) and polyclonal rabbit anti-human ADAMTS16 (#TA322059, AMS Biotechnology) (diluted 1:100 and 1:200, respectively, and incubated at room temperature for 1 h). Secondary antibodies were incubated at room temperature for 30 min and DAB was developed. Slides were counterstained with haematoxylin and mounted, and images were acquired on an Axioscan microscope slide scanner (Zeiss).

For peroxidasin immunostaining, there were 151 non-cancerous lung cores that were available for evaluation (149 interpretable following staining) and 138 tumour cores, of which 119 were paired samples (Supplementary Table 3). For ADAMTS16 immunostaining, there were 150 available non-cancerous lung cores (148 interpretable for inflammatory cells and 149 for non-tumour cells) and 155 available tumour cores (154 interpretable for tumour stroma and 152 for tumour cells), of which 133 were paired samples (Supplementary Table 3). Slides were scored for intensity of staining of tumour cells or stromal areas on a four-point scale: 0, no staining; 1, low staining; 2, moderate staining; 3, high staining. Data are presented as the proportion of positive staining within each category of staining for tumour and normal samples. Significant difference in the distribution of staining scores was determined by a chi-square test. Outcome data were recorded for each patient who had a tumour resection, with the median time to follow-up of 1,432 days (range 1,054–1,905 days).

Outcome included a record of death as defined by the clinical care team. Immunohistochemistry (IHC) scoring was assessed against survival for tumour staining.

### Differential gene expression analysis

Gene expression data were derived from RNA-seq datasets of primary tissue samples from cohorts of lung adenocarcinoma or lung squamous cell carcinoma patients extracted from TCGA, the Genotype-Tissue Expression repository and the Therapeutically Applicable Research to Generate Effective Treatments database using TNMplot (Bartha and Győrffy, 2021). Data from tumour samples were compared to paired data from matched adjacent normal samples.

### Survival analysis

Kaplan–Meier curves were computed from RNA-seq datasets of primary tissue samples from cohorts of lung adenocarcinoma or lung squamous cell carcinoma patients extracted from TCGA using KMplot (Lánczky and Győrffy, 2021). The lower and upper quartiles of gene expression were compared. Follow-up time was truncated to retain at least 10 patients at risk.

## Acknowledgements

We thank Jayne Culley, Kenneth Macleod and Emma Scholefield for technical assistance, Annya Smyth for help with tissue governance and Christopher Gregory for useful discussions. H.T. was funded by the EPSRC and MRC Centre for Doctoral Training in Optical Medical Imaging (EP/L016559/1). A.v.K. was supported by a Wellcome Trust multiuser equipment grant (208402/Z/17/Z). M.C.F. was funded by Cancer Research UK (C157/A15703 and C157/A24837). A.R.A. is supported by a Cancer Research UK Clinician Scientist Fellowship (A24867).

