## Supplementary Information for "Quantitative proteomics identifies tumour matrisome signatures in patients with non-small cell lung cancer"

Supplementary Figure 1

Supplementary Figure 2

Supplementary Figure 3

Supplementary Figure 4

Supplementary Table 1

Supplementary Table 2

Supplementary Table 3

[1]EPSRC and MRC CDT in Optical Medical Imaging, Queen's Medical Research Institute, University of Edinburgh, Edinburgh Bioquarter, Edinburgh EH16 4TJ, UK. [2]Centre for Inflammation Research, Queen's Medical Research Institute, University of Edinburgh, Edinburgh Bioquarter, Edinburgh EH16 4TJ, UK. [3]Cancer Research UK Scotland Centre, Institute of Genetics and Cancer, University of Edinburgh, Edinburgh EH4 2XR, UK. [4]Department of Pathology, Royal Infirmary of Edinburgh, Edinburgh EH16 4SA, UK. [5]Division of Molecular and Cellular Function, School of Biological Sciences, Faculty of Biology, Medicine and Health, University of Manchester, Manchester Academic Health Science Centre, Manchester M13 9PT, UK. [6]These authors jointly supervised the work. *Correspondence should be addressed to A.B. (; Twitter: @adambyron) or A.R.A. (; Twitter: @ahsanakram)

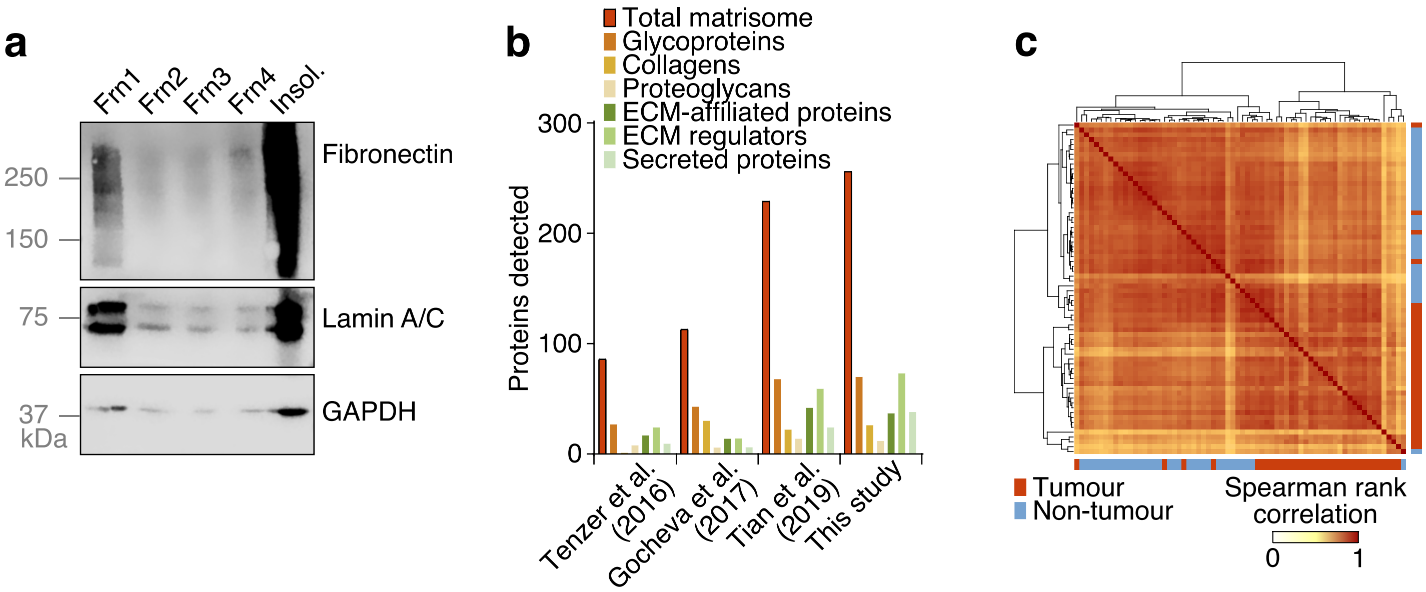


**Supplementary Fig. 1  Enrichment of ECM proteins from patient-derived lung tumours. a** Serially extracted fractions (Frn1–Frn4) were compared to the final insoluble ECM pellet (insol.) by western blotting to assess extraction efficacy of the ECM protein fibronectin. **b** Numbers of matrisome proteins detected in the current study compared to other recently published proteomic analyses of lung tumour tissue (details provided in Supplementary Table 2). **c** Correlation analysis of lung tissue samples. Spearman rank correlation coefficients for all pairwise sample comparisons were subjected to hierarchical clustering.

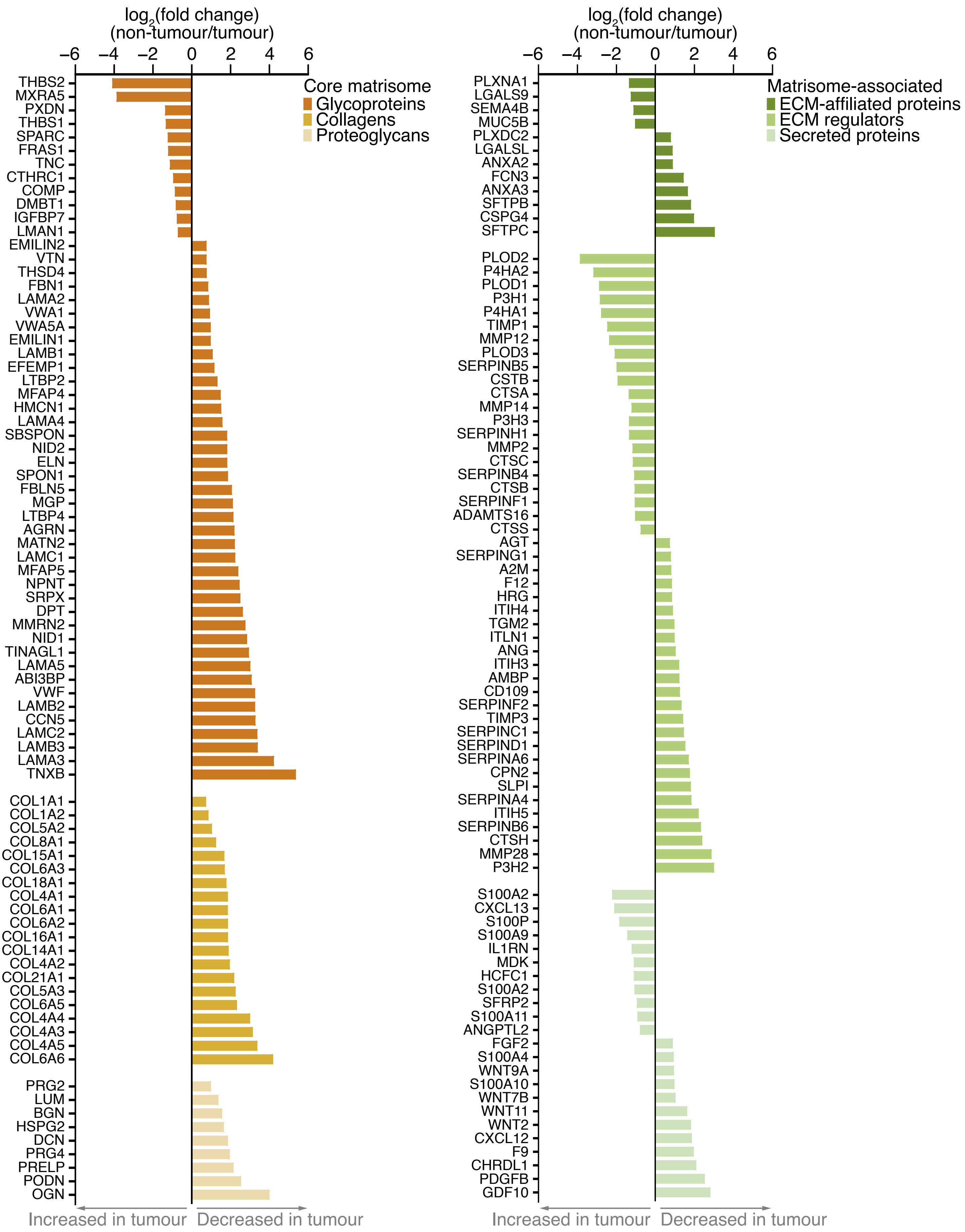


**Supplementary Fig. 2 Differential regulation of matrisome proteins between tumour and non-tumour samples.** Charts display $\log_2$-transformed label-free quantification intensity ratios (non-tumour/tumour) for differentially expressed core matrisome proteins (left panel) and matrisome-associated proteins (right panel) ($P < 0.05$, paired two-sided Student's $t$-test with Benjamini–Hochberg correction).

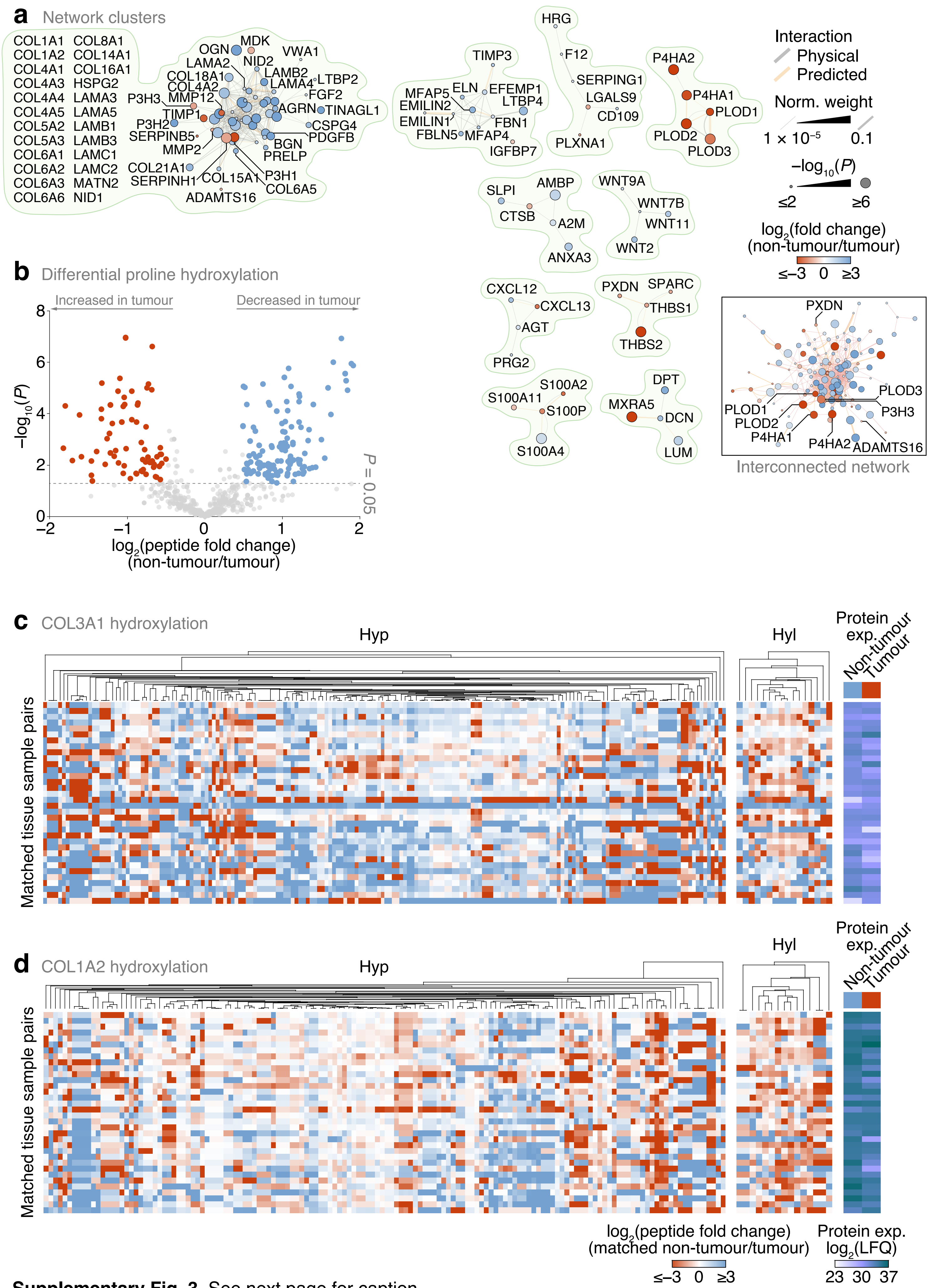


**Supplementary Fig. 3** See next page for caption.

**Supplementary Fig. 3 Regulation of proline and lysine hydroxylation in patient-derived lung tumour ECM.** **a** Markov clustering of the interaction network of proteins differentially regulated in tumour and non-tumour ECM ($P < 0.05$, paired two-sided Student's *t*-test with Benjamini–Hochberg correction). Proteins (nodes) are coloured according to enrichment or depletion in tumour samples and sized according to statistical significance. Protein interactions (edges) were weighted according to evidence of co-functionality. Clusters with at least four proteins are shown. Inset, complete connected network. **b** Volcano plot of peptides containing hydroxylated proline quantified by MS-based proteomics. Differentially regulated peptides are indicated with large coloured circles (red, increased in tumour; blue, decreased in tumour) ($P < 0.05$, FDR 20%, paired two-sided Student's *t*-test with Benjamini–Hochberg correction). **c**, **d** Regulation of proline and lysine hydroxylation in type III collagen α1 chain (COL3A1) (**c**) and type I collagen α2 chain (COL1A2) (**d**) across 34 matched non-tumour–tumour paired tissue samples. Total protein expression determined by label-free quantification (LFQ) shown for corresponding samples.

---

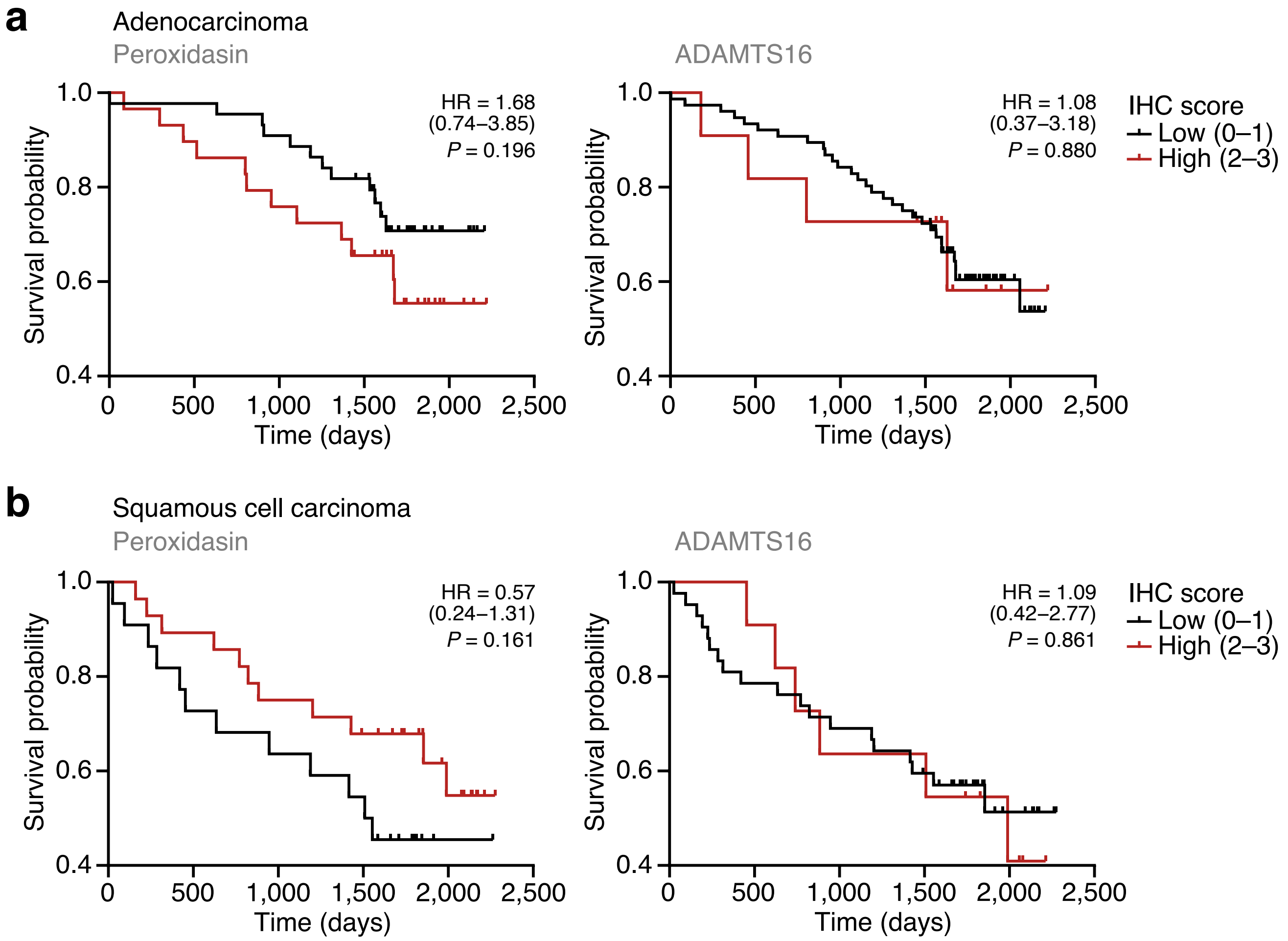


**Supplementary Fig. 4 Survival analyses for peroxidasin and ADAMTS16 immunostaining in the TMA cohort. a**, **b** Kaplan–Meier curves of patient survival associated with degree of tumour stroma immunostaining of peroxidasin (left panels) and ADAMTS16 (right panels) in lung adenocarcinoma (**a**) or lung squamous cell carcinoma (**b**) patients represented in the TMA cohort. Cores with uninterpretable immunostaining were omitted. HR, hazard ratio (with 95% confidence interval). Statistical analysis, log-rank test (*n* = 73 and 50 cores for adenocarcinoma and squamous cell carcinoma patients, respectively, for peroxidasin immunostaining and *n* = 87 and 53 cores for adenocarcinoma and squamous cell carcinoma patients, respectively, for ADAMTS16 immunostaining).

**Supplementary Table 1 Demographics of patients included in the study.**

| Characteristic | Category | Number of patients |
|---|---|---|
| Gender | Male | 14 |
| | Female | 21 |
| Age (years) | 40–49 | 1 |
| | 50–59 | 3 |
| | 60–69 | 10 |
| | 70–79 | 17 |
| | 80–89 | 3 |
| Tumour type | Adenocarcinoma | 17 |
| | Squamous cell carcinoma | 12 |
| | Large cell tumours | 3 |
| | Pleomorphic | 2 |
| Degree of tumour differentiation | Poor | 13 |
| | Moderate | 19 |
| | Well | 1 |
| | Not recorded | 1 |
| Tumour (T) stage | T1 | 1 |
| | T2 | 17 |
| | T3 | 10 |
| | T4 | 6 |
| Lymph node (N) stage | N0 | 24 |
| | N1 | 7 |
| | N2 | 3 |
| PET uptake | Low | 0 |
| | Moderate | 1 |
| | High | 32 |
| | Not recorded | 1 |
| Smoking status | Current | 9 |
| | Previous | 24 |
| | Never | 1 |
| Histology non-cancerous tissue | Normal | 7 |
| | Emphysema | 13 |
| | Emphysema and fibrosis | 1 |
| | Emphysema and inflammation | 1 |
| | Emphysema and pneumonia | 3 |
| | UIP fibrosis and emphysema | 1 |
| | Mild emphysema and non-caseating granuloma | 1 |
| | Inflammation | 1 |
| | Infract and pigment laden macrophages | 1 |
| | Pleural fibrosis | 1 |
| | Pneumonia | 1 |
| | Sarcoidosis | 1 |
| | Not recorded | 1 |

**Supplementary Table 2 Recently published proteomic analyses of lung tumour tissue against which our study was compared.**

| Study | Lung tissue source | Tissue extraction method | Proteolytic enzyme(s) | MS approach | Matrisome proteins identified |
|---|---|---|---|---|---|
| Tenzer et al. (2016) | Twenty-one patient tumour and matched non-tumour samples from 11 patients with adenocarcinoma and 10 patients with squamous cell lung carcinoma | Whole tissue lysates concurrently used for RNA extraction | Trypsin | High-definition MS (HDMS) | 86 matrisome proteins were detected. These proteins comprised 27 glycoproteins, 1 collagen, 8 proteoglycans, 17 ECM-affiliated proteins, 24 ECM regulators and 9 secreted proteins |
| Gocheva et al. (2017) | Three KP mice, where tumour formation was stimulated using Cre-expressing adenoviruses. Tumours were graded as stage 3–4. In addition, samples were analysed from C57BL/6J mice with bleomycin-induced pulmonary fibrosis and normal lung tissues | Fractionated ECM proteins separated using a commercial subcellular compartment protein extraction kit | PNGase F, Lys-C, trypsin | Tandem mass tag (TMT) | 100 matrisome proteins were detected initially, and 113 matrisome proteins were detected following exclusion of abundant spectra. These proteins comprised 43 glycoproteins, 30 collagens, 6 proteoglycans, 14 ECM-affiliated proteins, 14 ECM regulators and 6 secreted proteins |
| Tian et al. (2019) | Twenty patient samples with idiopathic pulmonary fibrosis and 20 control samples taken from patients undergoing surgery for pulmonary nodules/tumours | Whole tissue | Trypsin | Isobaric tags for relative and absolute quantitation (iTRAQ) | 229 matrisome proteins were detected. These proteins comprised 68 glycoproteins, 22 collagens, 14 proteoglycans, 42 ECM-affiliated proteins, 59 ECM regulators and 24 secreted proteins |

**Supplementary Table 3 Patient-derived lung cores included in the tumour microarray.**

| Staining | Characteristic | Number of cores | Proportion of cores (%) |
|---|---|---|---|
| Peroxidasin | Non-tumour (total available) | 151 | – |
| | Non-tumour (interpretable) | 150 | 100 |
| | Tumour (total available) | 138 | – |
| | Tumour (interpretable) | 138[a] | 100 |
| | Adenocarcinoma | 73 | 52.9 |
| | Squamous cell carcinoma | 51 | 37.0 |
| | Adenosquamous carcinoma | 5 | 3.6 |
| | Large cell carcinoma (other) | 7 | 5.1 |
| | Mixed small cell and large cell carcinoma | 1 | 0.7 |
| | Pleomorphic carcinoma | 1 | 0.7 |
| | Non-tumour–tumour sample pairs | 119 | – |
| ADAMTS16 | Non-tumour (total available) | 150 | – |
| | Non-tumour (interpretable) | 149 | 100 |
| | Tumour (total available) | 155 | – |
| | Tumour (interpretable) | 154[b] | 100 |
| | Adenocarcinoma | 87 | 56.5 |
| | Squamous cell carcinoma | 53 | 34.4 |
| | Adenosquamous carcinoma | 5 | 3.3 |
| | Large cell carcinoma (other) | 7 | 4.5 |
| | Mixed small cell and large cell carcinoma | 1 | 0.6 |
| | Pleomorphic carcinoma | 1 | 0.6 |
| | Non-tumour–tumour sample pairs | 133 | – |

[a]For tumour stroma only, 137 cores were interpretable

[b]For tumour cells only, 152 cores were interpretable
